# DHHC21 is a STIM1 protein S-acyltransferase that modulates immune function *in vivo*

**DOI:** 10.1101/2025.10.09.681383

**Authors:** Goutham Kodakandla, Ying Fan, Michael X Zhu, Savannah J. West, Askar Akimzhanov, Darren Boehning

## Abstract

Depletion of calcium from ER stores leads to the activation of calcium channels on the plasma membrane. This process is termed store-operated calcium entry (SOCE). The proteins STIM1 and STIM2 function as ER calcium sensors, and upon store depletion, they undergo a conformational change that allows them to bind to and gate Orai calcium channels on the plasma membrane. We have shown that both Orai1 and STIM1 are dynamically S-acylated after store depletion, which is required for SOCE. These results suggest the requirement of a calcium-activated protein S-acyltransferase such as DHHC21. Here, we show that DHHC21 is essential for SOCE *in vitro* and *in vivo*. Using the *depilated* mouse model that expresses DHHC21 but can no longer be activated by calcium, we show that DHHC21 activation is required for STIM1 S-acylation and subsequent calcium entry. Plasma membrane-localized DHHC21 is dynamically recruited into Orai1/STIM1 puncta upon store depletion, where it physically binds to STIM1. Finally, we show that *depilated* mice phenocopy many aspects of autoimmune lymphoproliferative syndrome (ALPS), including defective Fas-mediated calcium release, T cell death, neutropenia, and increased serum vitamin B12 levels. These results suggest that dynamic S-acylation has underappreciated and expansive roles in second messenger signaling and immune system function. Targeting DHHC21 may be therapeutically beneficial for ALPS and diseases associated with deregulated activation of STIM1, such as tubular aggregate myopathy and Stormorken syndrome.

## Introduction

Intracellular calcium levels regulate many physiological functions. Store-operated calcium entry (SOCE) is a mechanism where the depletion of endoplasmic reticulum (ER) calcium stores leads to the activation of calcium influx from the extracellular milieu (1-7). The ER calcium sensor Stromal Interaction Molecule 1 (STIM1) is activated upon depletion of calcium stores in the ER lumen, which undergoes a conformational change to approach the plasma membrane (PM), where it binds to and activates a calcium channel, Orai1 (8-11). This intermolecular complex between STIM1 and Orai1 is termed the calcium release-activated calcium (CRAC) channel (6, 12, 13). CRAC channels form macromolecular complexes near ER/PM junctions, which constitute the nexus of SOCE. These complexes have been described as ‘puncta’ owing to the punctate appearance in microscopic images (14). Abnormalities in CRAC channel function lead to disorders in the immune and musculoskeletal systems, such as severe combined immunodeficiency (SCID), tubular aggregate myopathy, and Stormorken syndrome (13, 15-19).

The mechanisms by which STIM1 and Orai1 proteins colocalize near ER:PM subdomains upon store-depletion have been a long-standing debate in the field. One prominent hypothesis is the diffusion trap model, which postulates a stochastic binding of activated STIM1 to Orai1 in these subdomains, where STIM1 ‘traps’ Orai1 to induce SOCE (20, 21). However, this model does not explain the active targeting of STIM1 to phosphatidylinositol 4,5-bisphosphate (PIP2) domains in membrane lipid rafts (22). In addition, studies have also shown that both Orai1 and STIM1 exist in dynamic equilibrium, where these proteins can transfer between different puncta (21). We have shown that S-acylation of Orai1 and STIM1 regulates CRAC channel formation (23-25). S-acylation is the reversible addition of lipid moieties to intracellular cysteine residues of target proteins (26). S-acylation is mediated by a set of enzymes known as palmitoyl acyltransferases (PATs). PATs are also known as DHHC enzymes owing to the presence of an aspartate-histidine-histidine-cysteine motif in the active site of these enzymes (27, 28). S-acylation affects protein stability, trafficking to subcellular compartments, and shuttling between membrane subdomains (29-33). We have shown that S-acylation actively targets Orai1 and STIM1 to puncta, providing an additional level of regulation to SOCE (23, 24). Our findings warrant refinement of the diffusion trap model.

Several DHHC enzymes are active in the PM, including DHHC5, DHHC20, and DHHC21 (34-37). We have shown that DHHC21 is essential for T cell calcium signaling by S-acylating components of both the Fas and the T cell receptor (TCR) complexes (36, 38). We also exploited the mouse model *depilated*, which has an in-frame deletion of a single phenylalanine residue in DHHC21 that eliminates a calmodulin-binding site (36, 39). *Depilated* mice have significant deficits in T cell signaling and differentiation into effector T cell lineages. *In vitro*, T cells from *depilated* mice have defective TCR signaling, including impaired activation of Lck, PLC-γ1, ZAP-70, and ERK pathways after TCR ligation (39, 40). Calcium transients induced by TCR ligation are significantly reduced in T cells from *depilated* mice, likely due to decreased S-acylation and activation of proteins such as Lck (36). The full repertoire of T cell signaling proteins regulated by DHHC21 S-acylation remains to be elucidated.

Based upon our previous findings in *depilated* mice, in this study, we examined the hypothesis that DHHC21 is the PAT for STIM1. We show that STIM1 S-acylation is significantly abrogated in spleens from homozygous *depilated* mice. This is associated with defects in splenocyte SOCE. We also show that DHHC21 is recruited into STIM1 puncta, and this is altered in *depilated* splenocytes. Consistent with the requirement of DHHC21 for Fas-mediated calcium release and cell death of T cells, we show that *depilated* mice phenocopy autoimmune lymphoproliferative syndrome (ALPS). We conclude that DHHC21 is the primary PAT for STIM1 *in vitro* and *in vivo*, and that *depilated* mice are a possible model for ALPS.

## Material and Methods

### Cells, antibodies, and constructs

HEK293T WT cells were purchased from American Type Culture Collection (ATCC) and cultured in Dulbecco’s Modified Eagle Medium (DMEM) supplemented with 10% fetal bovine serum (FBS), 1% L-glutamine, and 1% penicillin-streptomycin. DHHC21 knockout cells were generated using CRISPR-Cas9 plasmids obtained from Genscript Inc. The sgRNA sequences used in these plasmids for knockout of DHHC21 are 5’ AAGTGGTAGGGAACTCGCAG 3’, 5’ ATGAGACTAGCAGCCTTTAT 3’. Knockout clones were isolated by GFP selection followed by serial dilution to isolate isogenic clones. Knockout of DHHC21 was confirmed by Western blot. For experimental manipulations, cells were plated on polystyrene tissue culture dishes or 6-well plates. For imaging, cells were plated on poly-L-lysine-coated coverslips in 6-well plates. Cells were transfected with 0.5 µg STIM1 and 0.5 µg DHHC21 plasmids per 35-mm dish. All cells were maintained at 37ºC and 5% CO2 until use. Antibodies for immunoblotting and immunofluorescence imaging were purchased from commercially available sources: STIM1 (catalog no.: 4961S), calnexin (catalog no.: 2679), anti-rabbit immunoglobulin, horseradish peroxidase–linked secondary antibody (catalog no.: 7074S), anti-mouse immunoglobulin, and horseradish peroxidase–linked secondary antibody (catalog no.: 7076P2) were from Cell Signaling Technology; anti-DHHC21 (PA525096) was from Thermo Fisher Scientific; anti-CD3 human antibody (catalog no.: 14-0037-82) was from eBiosciences. JO2 antibody (catalog no.: 554255) was from BD Biosciences. The secondary antibodies for super-resolution STED imaging were obtained from Abberior Inc (STRED-1001, STRGREEN-1001). DHHC21-eGFP plasmids were purchased from Genscript Inc. The DHHC21-GFP plasmid was constructed by VectorBuilder. The DHHC21-FLAG plasmid was a kind gift from Dr. Masaki Fukata (National Institute for Physiological Sciences, Japan). The DHHC21-F233 mutant was generated using q5 site-directed mutagenesis kit from New England Biolabs using primers (F: TCAGAAGTTTTTGGCACTCGTTG, R: GGTCTGCTGCCATGGCTT). Mutagenesis was confirmed using Sanger Sequencing (Eton Biosciences). STIM1-mRFP was a generous gift from David Holowka and Barbara Baird (Cornell University). Thiol-Sepharose beads used for acyl-RAC were obtained from Nanocs, Inc and activated according to the manufacturer’s protocol before continuing with acyl-RAC protocol listed later. All other chemicals and reagents were purchased from Sigma–Aldrich or VWR.

### Mice

Wild type and *depilated* (zdhhc21^dep^) mice in the C56BL/6 background were bred in our barrier vivarium under pathogen-free conditions in accordance with the recommendations in the Guide for the Care and Use of Laboratory Animals of the National Institutes of Health. The animals were handled according to the animal care protocol #2020-1252 approved by the Rowan University Institutional Animal Care and Use Committee. Homozygous *depilated* mice had a higher mortality rate before the age of four weeks. To decrease the mortality rate, DietGel Boost was provided after birth as a high-calorie dietary supplement to the pups until they were weaned at 4 weeks of age. A qPCR strategy was used to genotype pups. *Depilated* mice were also confirmed visually by the loss of hair. Neutrophil counts were determined using a Zoetis Diagnostics Vetscan HM5 in male and female mice ranging from 4-52 weeks old. Mice used for other experiments were males and females, ranging in age from 6 to 8 weeks.

### Acyl-RAC assay

Acyl-RAC assay was performed as previously described by our group (41). Cell and tissue lysates were collected in lysis buffer (1% dodecyl-ß-D-maltoside in DPBS, supplemented with cOmplete protease inhibitor cocktail (Roche), 10 µM ML211 (acyl protein thioesterase inhibitor), and 10 mM phenyl-methyl-sulfonyl-fluoride (PMSF)). The lysates were centrifuged at 4 °C for 30 minutes at 20,000 xg. 500 µg of protein lysate were used for enrichment of the S-acylated proteome. Protein precipitation was performed using 2:1 methanol:chloroform, and the protein pellets were incubated with 0.1% methylmethanethiosulfonate (MMTS) for 20 minutes at 42 °C. Excess MMTS in the protein lysates was removed by three rounds of protein precipitation using methanol and chloroform. The resulting protein pellets were dissolved in 2SHB buffer (2% SDS, 5 mM EDTA, 100 mM HEPES, pH 7.4). 5% of the final volume was saved for input, and the remaining was used for acyl-RAC. 400 mM hydroxylamine (HA) was added to protein lysates to cleave the thioester bonds at pH 7. The lysates were incubated with thiol-Sepharose resin with rotation overnight at 4 °C. The next day, the samples were washed using wash buffer (1% SDS, 5 mM EDTA, 100 mM HEPES, pH 7.4). Proteins were eluted using 10 mM DTT in SDS buffer (1% SDS, 50 mM Tris-HCl, 10% glycerol, and 1% bromophenol blue) at 80 °C for 15 minutes with continuous shaking. Samples were then resolved on 10% SDS-PAGE gels and analyzed using Western blotting.

### Co-immunoprecipitation

Protein lysates were obtained from the spleens of WT and *depilated* mice using the lysis buffer used in the acyl-RAC section above. Protein lysates (1 mg) were incubated with 3 µl of either anti-DHHC21 or anti-FLAG antibodies overnight at 4 °C. The next day, 50 µl protein A agarose slurry (Pierce) was added to the samples, and the mixture was incubated at room temperature (RT). After one hour, samples were washed using the lysis buffer. After three washes, the agarose pellet was allowed to dry and subsequently quenched by boiling the samples with SDS buffer at 95 °C for 5 minutes. Samples were loaded on 10% SDS-PAGE gels and analyzed using western blotting.

### Super-resolution imaging

We performed STED super-resolution imaging using a STEDYCON microscope (Abberior). Splenocytes from *depilated* and WT mice were obtained, and immunofluorescence was performed on these cells as previously described elsewhere (42). Briefly, the cells were plated on poly-L-lysine coated coverslips and treated with anti-CD3 where required. After treatment, the cells were fixed using 4% ice-cold paraformaldehyde for 20 minutes at RT with rotation. The cells were quenched with 30 mM glycine/PBS solution for 5 minutes at RT with rotation. Next, the cells were washed 3 times with PBS with rotation. Cells were permeabilized with 0.25% Triton X-100 and 1% BSA in PBS for 10 minutes at RT with rotation. Cells were then incubated with 2% BSA in PBS for 1 hour at RT with rotation. Cells were washed 3 times with PBS for 5 minutes each and then incubated with antibodies in PBS with 0.3% BSA. Cells were washed and mounted on glass slides sealed with nail polish. The secondary antibodies for STED imaging were Star Red and Star Green anti-rabbit or anti-mouse obtained from Abberior Inc.

### Fura-2 imaging

Fura-2 imaging on WT HEK293, DHHC21-KO HEK293, and mouse splenocytes was performed as previously described (23, 24). Briefly, for adherent cells, 2 days before imaging, cells were seeded on poly-lysine coated coverslips in 6-well tissue culture plates. The next day, these cells were transfected with DHHC21 and STIM1 plasmids. Imaging buffer contained 0.1% bovine serum albumin, 107 mM NaCl, 20 mM HEPES, 2.5 mM MgCl_2_, 7.5 mM KCl, 11.5 mM glucose, and 1 mM CaCl_2_, pH 7.2. For store depletion, imaging buffer devoid of CaCl_2_ was used. 24h after transfection, the cells were incubated with 5 µM Fura-2 in the imaging buffer for 30 minutes at RT, followed by de-esterification for 30 minutes at RT. Time-lapse images were recorded on a Nikon Ti-2 microscope using a 40X oil immersion objective. Cells were excited alternatively at 340 nm and 380 nm every 2 seconds for 16 minutes, and the fluorescence emission was collected at 525 nm. The first minute of the time-lapse was used to obtain the baseline calcium level. After 1 minute, thapsigargin (TG, 10 µM) in the calcium-free imaging buffer was added to induce store depletion. After an additional 7 minutes, the buffer was replaced with calcium-replete imaging buffer with TG to measure calcium entry. The ratio of fluorescence at 340 to 380 nm was used to quantify cellular calcium levels. Cells that did not respond to TG were excluded from our analysis. Identical exposure parameters were used to obtain images. Peak entry was calculated by subtracting the maximum fluorescence ratio (Rmax) after calcium addback from the fluorescence ratio at time 0 (R_0_) and normalized to R_0_.

### Total internal reflection fluorescence imaging (TIRF)

Cells were seeded on poly-L-lysine-coated coverslips 2 days before TIRF imaging. Cells were transfected with WT or F233 DHHC21-eGFP, WT STIM1-mRFP, and WT Orai1-Myc using Lipofectamine 3000 following the manufacturer’s protocol. A Nikon Eclipse Ti microscope, equipped with a TIRF illumination system, was used for imaging. Coverslips were mounted in Attofluor chambers with the imaging buffer containing calcium for live imaging using a 60x oil-immersion TIRF objective. Cells were alternatively excited at 488 nm and 561 nm lasers every 5 seconds. The first minute of the recording was used to obtain baseline colocalization levels. The images were captured over an 8-minute period. After 1 minute, the imaging buffer was replaced with 10 µM TG in the calcium-free imaging buffer. Cells that had puncta prior to store depletion or those that showed high colocalization in the resting state were excluded from analysis. Colocalization analysis was performed using Nikon NIS Elements software. Using regions of interest around cells, we obtained Pearson’s correlation coefficients between both channels for the entire time series. We normalized correlation coefficients to time zero by dividing the correlation coefficient at a given time (Rn) by time zero (R_0_). These values were used to plot the colocalization time curve. Peak colocalization was calculated by subtracting peak colocalization after TG addition (Rmax) from colocalization just before TG addition (R_60_) and dividing the difference by R_60_. ((Rmax-R_60_)/R_60_).

### Confocal imaging

DHHC21-KO cells plated on coverslips and transfected with WT and F233 versions of DHHC21-eGFP were used for confocal imaging. The cells were imaged using the confocal mode on the STEDYCON microscope described above, using a 100x oil objective. Identical exposure parameters were used to obtain images from WT and F233 DHHC21 expressing cells.

### Caspase-3 activity assay

Splenocytes from WT or homozygous zdhhc21^dep/dep^ mice were treated with JO-2 for 24h. Total caspase activity was determined in cell lysates using the caspase-3 substrate Z-DEVD-R110 as described previously (43). The production of fluorescent substrate was monitored continuously every minute for 1h in a microplate reader. The slope of the linear regression drawn through each time point was used to determine the change in fluorescence over time for each sample.

### ELISA

ELISA tests for ANA, CRP, and VB12 were purchased from ThermoFisher (501487635, EM20RB, 502287196). Serum was collected from 6-8-week-old mice. The total amounts of each analyte were determined as per the manufacturer’s instructions.

## Results

### DHHC21 is required for STIM1 S-acylation

We have previously shown that CRAC channel components Orai1 and STIM1 are S-acylated (23, 24). *Depilated* mice have an in-frame deletion of phenylalanine 233 (F233) residue in the C-terminal tail of DHHC21 that eliminates calmodulin binding (36, 39). This mutation leads to deficits in TCR signaling and T cell differentiation. We have concluded that the ΔF233 mutation is likely a loss-of-function mutation (36, 39). To determine if *depilated* mutant mice have deficits in STIM1 S-acylation *in vivo*, we harvested spleens from wild-type and *depilated* mice and performed an acyl-RAC assay to purify S-acylated proteins. We found that STIM1 has significantly reduced S-acylation in *depilated* spleen (Figure 1A,B). The S-acylation of DHHC21 and the ER protein calnexin was unchanged. Total levels of all three proteins were similar in wild-type and homozygous mutant mice. Overexpression of wild-type or ΔF233 DHHC21 in HEK cells expressing endogenous DHHC21 did not significantly affect S-acylation of STIM1, indicating that the ΔF233 mutant enzyme does not function as a dominant-negative inhibitor of the wild-type protein (Figure 1C). We next determined if SOCE was altered in WT and homozygous *depilated* splenocytes. Resting (baseline) calcium levels were significantly higher in *depilated* homozygous splenocytes, indicating significant alterations in calcium homeostasis (Figures 1D,E). To measure SOCE, we depleted ER stores with thapsigargin (TG) in a calcium-free solution and then added back calcium to measure entry (Figure 1D). We found that homozygous *depilated* splenocytes had significantly reduced SOCE (Figure 1F) and a smaller TG-releasable pool (Figure 1G). These results indicate DHHC21 is the primary PAT for STIM1 *in vitro* and *in vivo*, and is necessary for efficient SOCE.

**Figure 1:**
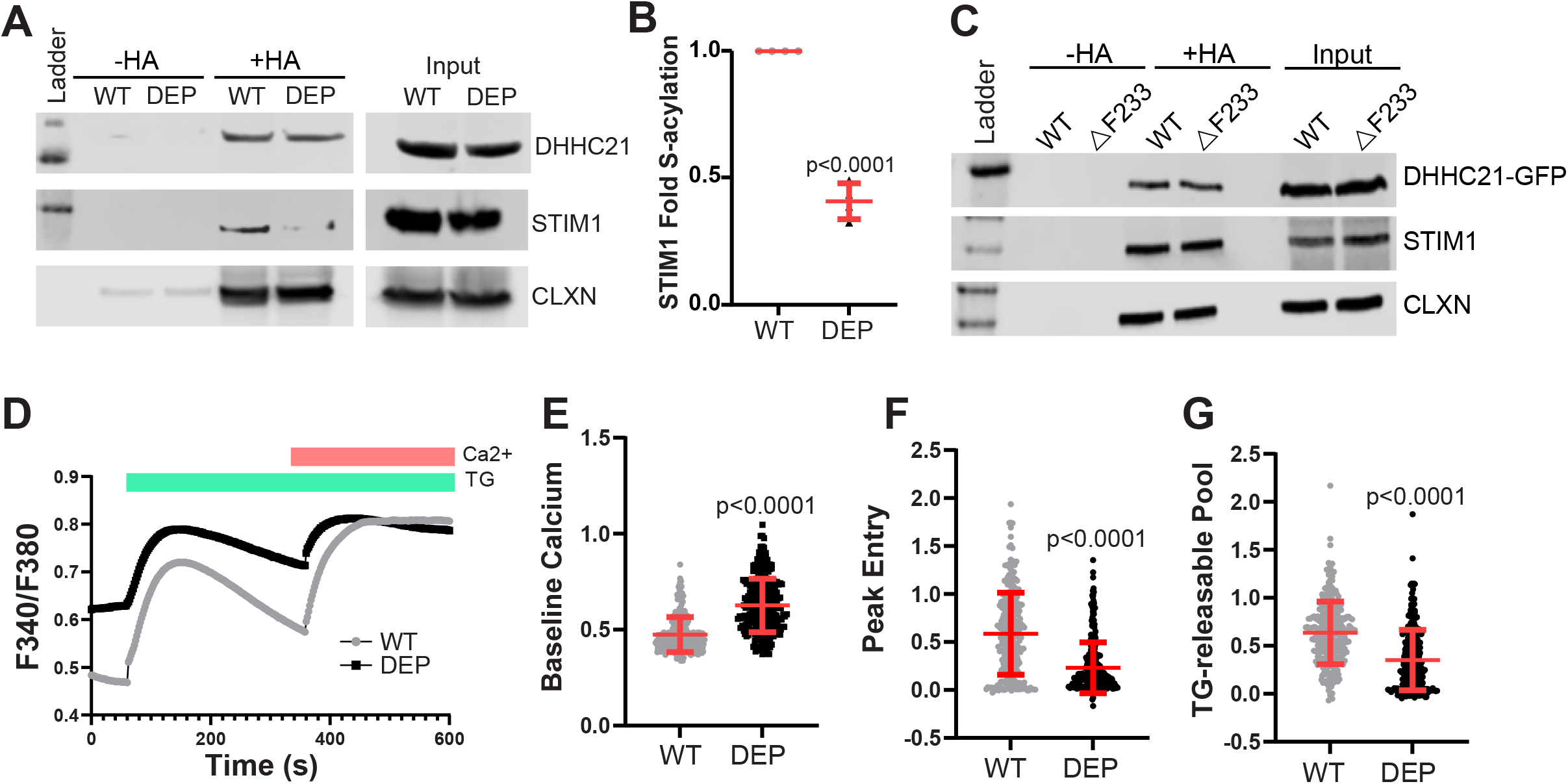
DHHC21 is the STIM1 protein acyltransferase and regulates store-operated calcium entry. A) Spleens from C57BL6F1/J WT and *depilated* (DEP) homozygous mice were collected, and acyl-RAC was performed. Reactions without hydroxylamine (-HA) are a negative control. Calnexin was used as both a positive control and a loading control for S-acylated proteins. B) Quantification of fold S-acylation of STIM1 normalized to calnexin from four independent experiments. Statistical significance between the groups was determined using Student’s t-test. Error bars indicate S.D. C) WT and ΔF233 DHHC21-GFP plasmids were co-expressed along with WT STIM1-mRFP and WT Orai1-Myc in DHHC21-KO cells, followed by acyl-RAC. D-G) Splenocytes were collected from C57BL6 WT and *depilated* homozygous mice and used for Fura2 imaging using 10 µM thapsigargin (TG) for store-depletion and 1 mM calcium addback to assess store-operated calcium entry. F340/F380 ratios were used to quantify calcium levels in these cells. Representative traces are shown in (D). Baseline calcium levels (E), peak entry (F), and TG-releasable pool (G) were calculated from the F340/F380 values. Statistical significance between the groups was determined using Student’s t-test. Error bars indicate S.D.

### DHHC21 is required for store-operated calcium entry

To determine the role of DHHC21 in SOCE, we generated a DHHC21 CRISPR knockout HEK293 cell line. We confirmed a complete loss of expression by Western blotting (Figure 2A). We found that DHHC21 knockout cells had significantly reduced SOCE induced by TG (Figures 2B,C). We tried to rescue DHHC21 KO cells by transfecting the cells with WT and ΔF233 DHHC21. While expression of WT DHHC21 completely rescued SOCE in DHHC21 KO cells, that of ΔF233 DHHC21 only partially restored SOCE, showing significantly less effective rescue than wild-type (Figure 2D, E). Interestingly, this *in vitro* rescue model with ΔF233 in a null DHHC21 background recapitulated the increased baseline calcium levels found in *depilated* splenocytes (Figure 2F). This suggests that ΔF233 has direct effects on baseline calcium independent of a complete loss of function. Finally, we found that the TG-releasable pool was similar between WT and ΔF233 rescue (Figure 2G). We conclude that in this independent model, DHHC21 is required for SOCE, and the *depilated* ΔF233 mutation cannot fully rescue a cellular model with a complete loss of DHHC21 function.

**Figure 2:**
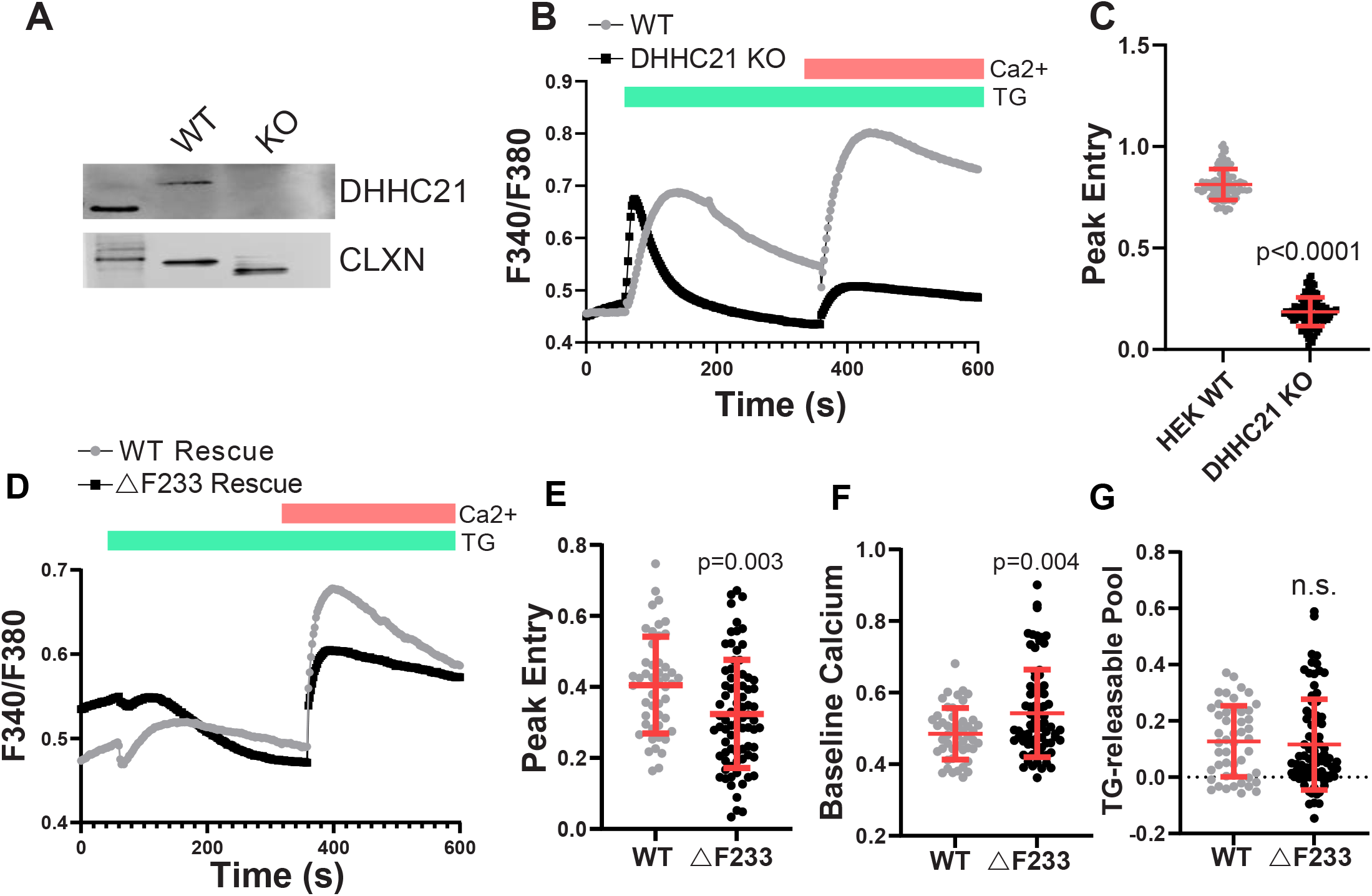
DHHC21 KO cells have impaired SOCE. Clonal DHHC21 CRISPR knockout HEK293 cells were generated as described in the Methods section. A) Western blotting was performed on whole cell lysates obtained from WT and DHHC21-KO cells and probed for DHHC21. Calnexin was used as a loading control. B) Fura2 imaging was performed on WT and DHHC21-KO cells. Store depletion was induced by 10 µM thapsigargin (TG) in calcium-free buffer. Calcium (1 mM) was added back after 6 minutes to allow calcium entry. C) Peak calcium entry in WT and DHHC21 KO cells. Statistical significance between the groups was determined using Student’s t-test. Error bars indicate S.D. D-G) Wild-type or ΔF233 DHHC21-eGFP was co-expressed with STIM1-mRFP and Orai1-Myc in DHHC21-KO cells, and Fura2 imaging was performed. Store-operated calcium entry was determined as in (B). Representative traces are shown in (D). Peak calcium entry (E), baseline calcium (F), and the TG-releasable calcium pool (G) were quantified for WT and ΔF233 cells. Statistical significance between the groups was determined using Student’s t-test. Error bars indicate S.D.

### Co-localization of WT and ΔF233 DHHC21 with STIM1

If DHHC21 is a protein S-acyltransferase for STIM1, the two proteins should interact (at least transiently). We made lysates from the spleens of WT and homozygous *depilated* mice, and co-immunoprecipitated STIM1 with DHHC21. Interestingly, STIM1 binds avidly to DHHC21 in spleens from both WT and *depilated* mice, exhibiting even increased binding to ΔF233 DHHC21 in the mutant samples (Figure 3A). We hypothesized that binding of DHHC21 to STIM1 may increase after store depletion. To test this hypothesis, we performed TIRF imaging of DHHC21-GFP and STIM1-mRFP in DHHC21 KO cells. As shown in Figures 3B-D and Supplementary Movies 1 and 2, store depletion leads to rapid recruitment of both WT and ΔF233 DHHC21 into STIM1 puncta. Consistent with the co-immunoprecipitation results, we observed a significantly higher peak colocalization of ΔF233 DHHC21 compared to WT DHHC21. We confirmed that the endogenous DHHC21 and STIM1 proteins in HEK293 cells bind after store depletion using superresolution imaging (Supplementary Figure 1). To determine the recruitment of DHHC21 to STIM1 in splenocytes, we performed superresolution imaging on splenocytes obtained from WT and *depilated* mice. We treated splenocytes *in vitro* with anti-CD3 antibody to activate TCR signaling and SOCE prior to fixation and staining for DHHC21 and STIM1. As shown in Figures 3E-F, we detected a significantly higher colocalization between DHHC21 and STIM1 in splenocytes treated with anti-CD3. The co-localization of DHHC21 and STIM1 in *depilated* mice was even more pronounced, corroborating our findings by TIRF imaging in the in vitro model. Together, these results indicate that ΔF233 DHHC21 binds STIM1 more avidly than the WT protein. We interpret these findings to mean that ΔF233 DHHC21 binds STIM1 but does not release the protein because it cannot complete the enzymatic cycle. These results also suggest that calcium-calmodulin binding to DHHC21 regulates enzymatic activity independently of substrate binding.

**Figure 3:**
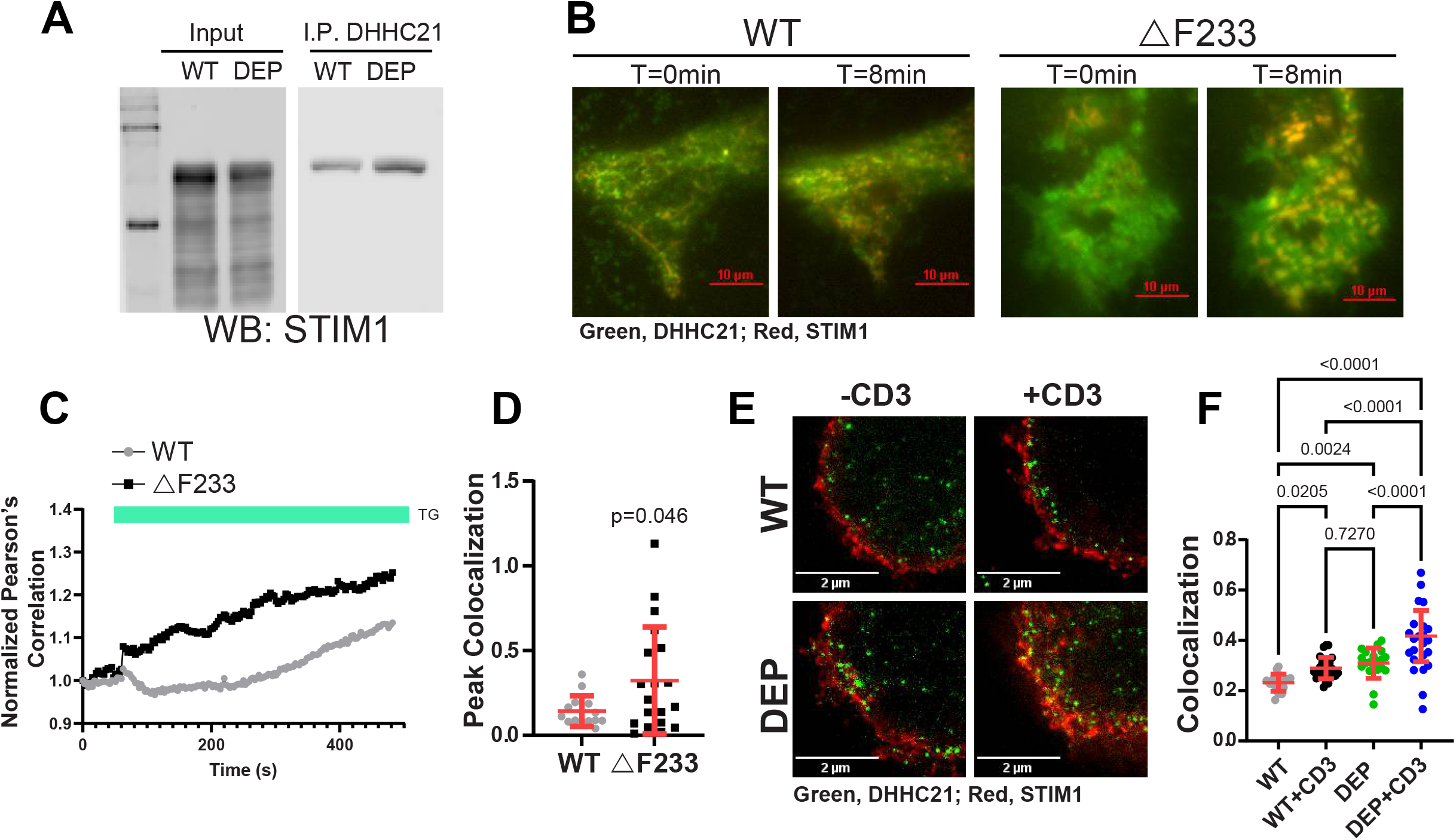
Store depletion increased colocalization of DHHC21 with STIM1. A) Co-immunoprecipitation of STIM1 by DHHC21 from WT and *depilated* (DEP) splenocytes. B) DHHC21-KO HEK293 cells were transfected with WT or ΔF233 DHHC21-eGFP, STIM1-mRFP, and Orai1-Myc plasmids, and store-depletion was induced using 10 µM thapsigargin (TG). Representative merged images are shown for a single experiment at times zero and at eight minutes after TG treatment. (C-D) Peak colocalization was quantified using Pearson’s correlation values obtained from WT and ΔF233 expressing cells. Statistical significance between the groups was determined using Student’s t-test. Error bars in (D) indicate S.D. E) Splenocytes were obtained from C57BL6F1/J WT and *depilated* homozygous mice and used for STED imaging. Store depletion was induced by anti-CD3 antibodies. F) Pearson’s correlation between DHHC21 and STIM1 before and after store depletion. Significance was determined using a Student’s t-test. Error bars indicate S.D.

### *Depilated* mice phenocopy autoimmune lymphoproliferative syndrome (ALPS)

We have previously shown that Fas signaling in T lymphocytes requires calcium release (38, 44). Engagement of the calcium release machinery requires rapid and dynamic S-acylation of signaling proteins such as the tyrosine kinase Lck (44). Mutations in the Fas pathway cause ALPS in patients (45). The primary diagnostic criteria for ALPS are chronic lymphadenopathy and splenomegaly, increased peripheral CD4-/CD8-T cells, defective Fas-mediated lymphocyte apoptosis, and mutations in a Fas pathway gene (46). Secondary criteria include increased serum vitamin B12 levels, autoimmune cytopenia, and increased IgG levels. However, about 10-20% of patients have no known genetic mutation (ALPS-U), suggesting that additional unknown genes contribute to disease progression (47). We have previously shown that *depilated* mice have marked lymphadenopathy and splenomegaly (39). In addition, we discovered profound defects in T cell development in these mice (39). We have also shown, using shRNA-mediated knockdown, that DHHC21 is required for Fas-mediated calcium release (44). Thus, we hypothesized that *depilated* mice may phenocopy ALPS. *Depilated* mice demonstrate a failure to thrive as determined by weight in both young and adult mice (Figure 4A). We found that splenocytes isolated from homozygous *depilated* mice had significant defects in both Fas-mediated calcium release and apoptotic cell death as determined by caspase-3 enzymatic activity (Figures 4B-D). This was associated with significantly elevated serum vitamin B12 levels (Figure 4E). C-reactive protein (CRP), a marker of inflammation and autoimmunity, was also elevated in the sera of homozygous mutant mice (Figure 4F). Similarly, we found a significant elevation of the autoimmunity marker anti-nuclear antibody (ANA) in the sera of homozygous mice (Figure 4G). Finally, we found age-dependent neutropenia in homozygous mutant mice (Figure 4H). We conclude that *depilated* mice phenocopy ALPS and suggest that mutations in DHHC21 should be investigated in ALPS-U patients who do not have a mutation in a known causative gene.

**Figure 4:**
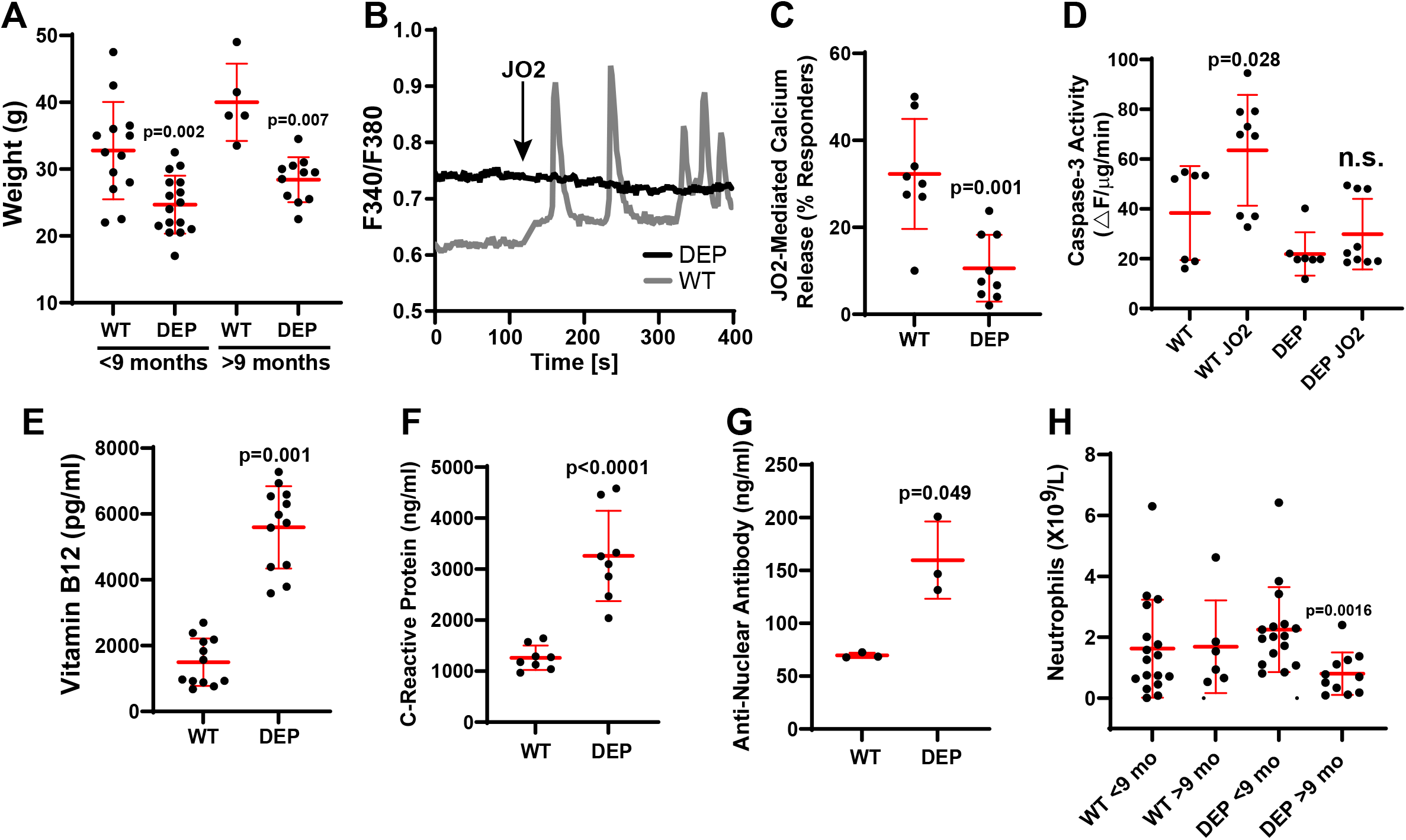
*Depilated* mice phenocopy ALPS. A) WT and *depilated* (DEP) mice younger and older than 9 months were weighed and analyzed using Student’s t-test. Error bars represent SD. B-C) Splenocytes from C57BL6F1/J WT and *depilated* homozygous mice were isolated, and Fura2 calcium imaging was performed. Representative splenocyte responses to stimulation with the Fas agonistic antibody JO2 (B) and quantification of the percent responders to JO2 averaged from eight (WT) or nine (*depilated*) mice (C) are shown. D) Caspase-3 enzymatic activity +/-JO2 in WT and *depilated* splenocytes. (E-G) Serum Vitamin B12 (E), C-reactive protein (F), and anti-nuclear antibody (G) levels in WT and *depilated* mice. H) Neutrophil counts in WT and *depilated* mice younger and older than 9 months. Data were analyzed using the Student’s t-test. Error bars represent SD.

## Discussion

S-acylation plays a crucial role in store-operated calcium entry by modifying the localization and activity of Orai1 and STIM1. Previously, we have shown that both Orai1 and STIM1 undergo S-acylation upon store depletion. S-acylation directs these proteins to membrane subdomains where they form CRAC channels to promote SOCE. Cysteine mutant versions of these proteins that cannot undergo S-acylation show deficits in SOCE. Previously, another group demonstrated that DHHC20 S-acylates Orai1 (48). In this work, we show that DHHC21 is the PAT that S-acylates STIM1 *in vivo* using *depilated* mice that have the ΔF233 mutation in DHHC21. We also show that ΔF233 DHHC21 can still bind but cannot S-acylate STIM1. This leads to significant deficits in SOCE and immune system function *in vivo*. It remains to be determined whether Orai1 can also be S-acylated by DHHC21, or whether two different PATs are required for SOCE after store depletion.

DHHC21 is a plasma membrane-localized protein S-acyltransferase. We have shown previously that DHHC21 plays a key role in S-acylation of many proteins involved in T cell receptor signaling. TCR components such as PLC-γ1, Lck, and ZAP-70 undergo S-acylation upon treatment with anti-CD3, which activates the T cell receptor (36). We have previously shown that Jurkat cells with DHHC21 knocked down by shRNA or CD4+ T cells isolated from *depilated* mice are defective in S-acylation of these proteins and downstream signaling events, indicating that DHHC21 plays a prominent role in TCR signaling (36, 39, 44). This formed the basis for our hypothesis that DHHC21 might mediate the S-acylation of STIM1. The S-acylation of STIM1 by PM-localized DHHC21 strongly suggests that S-acylation of the STIM1 tail stabilizes its PM association. The polybasic domain of STIM1 has been shown to interact with membrane phospholipids such as PI(4,5)P_2_ and physically bind to the PM. In addition, isoleucine 384 (I384) in the cholesterol-binding site within the SOAR/CAD domain of STIM1 has also been shown to bind to the PM upon store-depletion induced by thapsigargin. This interaction was shown to enhance SOCE. Together, these domains stabilize the active conformation of STIM1. The redundant C-terminal domains mediating PM binding of STIM1 may explain the partial SOCE retained in *depilated* splenocytes and DHHC21 KO HEK293 cells. Further experiments are warranted to determine the mechanism(s) by which SOCE is partially retained in *depilated* splenocytes.

We observed differences in SOCE in *depilated* splenocytes compared to CRISPR-KO HEK293 cells (Figures 1,2). In addition, rescue experiments demonstrated that ΔF233 DHHC21 expression has direct effects on baseline calcium in KO cells, indicating that this mutant enzyme has significant consequences for calcium homeostasis (Figure 2F). We conclude that ΔF233 DHHC21 is not a complete loss of function. Indeed, we found that this enzyme can bind STIM1 even more avidly than the WT enzyme (Figure 3). This was somewhat unexpected. We hypothesize that DHHC21-ΔF233 is still competent to bind substrate (including STIM1), but it can no longer release the substrate because of a lack of enzymatic activity. It is likely that ΔF233 DHHC21 sequesters additional substrates at the PM with significant ramifications for cellular physiology. Recently, a mouse model with a T-cell-specific knockout of DHHC21 was described (49). Like our previous studies on T cell development and differentiation in *depilated* mice (36, 39), the researchers found significant effects on the peripheral T cell compartment, but with many important differences. Perhaps most strikingly, they found that both spleen and lymph node sizes were smaller with fewer total cell counts in contrast to the splenomegaly and lymphadenopathy found in *depilated* mice. Thus, their findings support our hypothesis that ΔF233 DHHC21 is not a loss-of-function mutation and, indeed, may have significant gain-of-function effects on T cell signaling. It was found that 21 proteins in T cells are likely substrates of DHHC21 using proximity labelling (49), all of which could potentially be sequestered by ΔF233 DHHC21 to affect T cell function. Future work will confirm this hypothesis and further explore the ALPS-like phenotype in *depilated* mice.

## Acknowledgements

We thank David Holowka and Barbara Baird (Cornell University) for the gift of the STIM1-mRFP plasmid and Masaki Fukata (National Institute for Physiological Sciences, Japan) for the kind gift of the DHHC21-FLAG plasmid. We wish to thank the members of the Boehning and Akimzhanov labs for many helpful discussions.

## Data availability

All data supporting this study are available from the corresponding authors upon request.

---

**Supplementary Figure 1:**
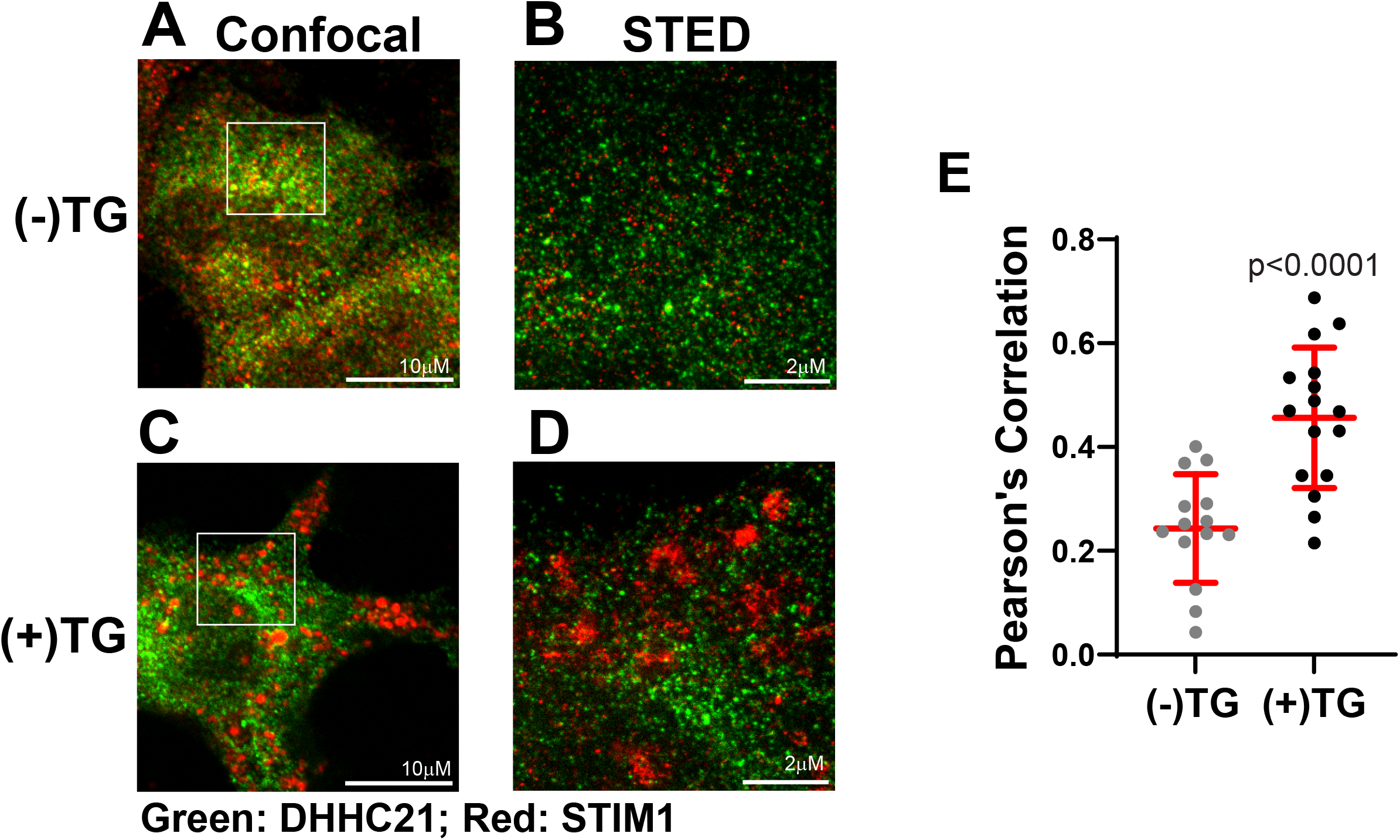
Endogenous DHHC21 colocalizes with STIM1 in HEK293 WT cells. Confocal (A and C) and STED (B and C) images of HEK293 WT cells untreated (A and B) or treated (C and D) with 10 μM thapsigargin (TG). The boxes in Panels A and C were used as the imaging area for STED imaging in Panels B and D. Green represents DHHC21, and red represents STIM1. E) Colocalization was quantified using Pearson’s correlation values obtained from cells. Statistical significance between the groups was calculated using Student’s t-test. Error bars indicate S.D. Pearson’s correlation between DHHC21 and STIM1 obtained from (E) using the JACoP plugin was analyzed using a Student’s t-test. Error bars indicate S.D.

**Supplementary Video 1: DHHC21 and STIM1 colocalize upon ER store depletion**. DHHC21 KO cells were transfected with DHHC21-GFP and STIM1-mRFP, and TIRF microscopy was performed. Store depletion was induced using 10 µM thapsigargin (TG).

**Supplementary Video 2: ΔF233 DHHC21 and STIM1 colocalize upon ER store depletion**. DHHC21 KO cells were transfected with ΔF233 DHHC21-GFP and STIM1-mRFP, and TIRF microscopy was performed. Store depletion was induced using 10 µM thapsigargin (TG).

